# HYPER-Net: Physics-Conditioned Self-Supervised Reconstruction for Fourier Light-Field Microscopy

**DOI:** 10.64898/2026.04.14.718527

**Authors:** Zhi Ling, Xuanwen Hua, Wenhao Liu, Hao Wu, Lucinda M Peng, Jessica Hou, Parvin Forghani, Christopher Pierce, Ge-Ah R Kim, Shuichi Takayama, Shuyi Nie, Chunhui Xu, Hang Lu, Shu Jia

## Abstract

The rapid convergence of optical innovation and machine intelligence is reshaping biological imaging by enabling platforms that jointly advance image formation and computational reconstruction for highspeed, high-resolution volumetric microscopy. However, broadly accessible three-dimensional imaging at high spatiotemporal resolution remains limited by the reliance of existing supervised methods on large modality-matched training datasets, the computational burden of conventional iterative reconstruction, and sensitivity to optical mismatch arising from small deviations in the spatial-angular point spread functions. Here, we introduce HYPER-Net, a physics-conditioned self-supervised framework for Fourier light-field microscopy that integrates scan-free volumetric acquisition with fast, robust three-dimensional reconstruction. HYPER-Net incorporates experiment-specific point-spread functions into the learning process in two complementary roles: as the forward operator that enforces measurement consistency and as a conditioning signal that adaptively modulates intermediate feature representations. This design reduces reliance on paired experimental ground-truth volumes, improves robustness to system variation, and enables generalizable reconstruction across diverse biological contexts. Using human colon organoids, embryonic *Xenopus* laevis hearts, hiPSC-derived cardiac spheroids, and freely moving *Caenorhabditis elegans*, we demonstrate high-fidelity volumetric imaging of tissue morphology, cardiac function, calcium-contraction coupling, and locomotion-associated neural and muscular dynamics. These results position HYPER-Net as a versatile framework for rapid volumetric imaging and quantitative analysis of dynamic biological systems across basic research and biomedical applications.

## INTRODUCTION

Microscopy is a foundational technology in the life sciences, enabling visualization of biological systems across scales, from single molecules to whole organisms^1,2^. Continued advances in imaging performance, including gains in spatial resolution, temporal speed, and experimental scale, have greatly expanded the scope of biological interrogation^3–5^. Meanwhile, these advances have also increased data dimensionality and analytical complexity, creating substantial demands for computational reconstruction, processing, and quantitative interpretation^6^. Artificial intelligence, particularly machine learning and deep learning, is increasingly transforming microscopy workflows, spanning data acquisition, image restoration, segmentation, and downstream biological analysis^7–9^. The convergence of optical innovation and machine intelligence has therefore become an emerging driver of progress in guided image acquisition^10^, denoising^11^, volumetric reconstruction^12^, segmentation and classification^13^, and predictive biological inference^14^.

In parallel, the demand for imaging fast biological dynamics has stimulated the development of light-field microscopy (LFM), which enables snapshot volumetric imaging at camera-limited speeds^15–18^. By using a microlens array to encode both spatial and angular information from the emitted fluorescence into a single two-dimensional measurement, LFM enables computational recovery of the full three-dimensional sample volume from a single exposure^15–17^. Fourier light-field microscopy (FLFM)^19–22^ further improves this framework by recording the light field at the Fourier plane, thereby providing axially uniform magnification and a laterally invariant point spread function (PSF). These properties improve reconstruction fidelity while reducing computational burden, facilitating integration with complementary optical and computational strategies, and broadening applicability across diverse biological systems^23–30^.

Despite these advantages, light-field reconstruction remains a major bottleneck. Conventional reconstruction typically relies on iterative deconvolution, such as Richardson–Lucy (RL), which is computationally intensive and often requires many iterations to reach acceptable image quality^31^. This limitation constrains throughput, particularly for long time-lapse experiments and large volumetric datasets. To address this challenge, several supervised learning approaches have recently been developed to accelerate volumetric inference by learning direct mappings from two-dimensional light-field measurements to three-dimensional volumes^32–36^. However, these methods generally rely on high-quality paired training data or sample-specific structural priors, which can limit their generalizability across specimens with distinct morphologies, labeling patterns, or cellular organization (**Supplementary Table 1**). This limitation becomes especially pronounced for organ- and tissue-level samples, where marker types, cell densities, and structural heterogeneity can vary substantially across experiments.

A further challenge arises from the sensitivity of light-field reconstruction to accurate PSF modeling. Because axial information is multiplexed into a two-dimensional projection, even slight microlens misalignment or system drift can alter the image formation process and degrade inverse reconstruction when previously measured or simulated PSFs are used. As a result, supervised models often require retraining after optical realignment or gradual system perturbation, especially in experiments conducted over extended periods^17,37^. Without such recalibration, a mismatch between modeled and experimental PSFs can lead to volumetric distortion, spatial misregistration, and loss of genuine biological information.

To overcome these challenges, we present HYPER-Net, a physics-conditioned, self-supervised framework for hybrid PSF-embedded reconstruction. Unlike prior self-supervised approaches, which enforce measurement consistency only through the forward projection of the reconstructed volume^12,37,38^, HYPER-Net explicitly incorporates experiment-specific PSFs into the learning process via feature-wise linear modulation (FiLM), allowing intermediate feature representations to be adaptively conditioned on the optical state of the imaging system. This strategy enables robust and generalizable reconstruction under optical perturbation and across biologically diverse datasets. HYPER-Net exhibits substantially greater robustness to system deviations than a self-supervised baseline without FiLM conditioning and outperforms conventional deconvolution in both reconstruction fidelity and computational efficiency. We demonstrate that HYPER-Net enables rapid, high-fidelity, and high-throughput volumetric imaging across multiple dynamic biological models, including human colon organoids, embryonic *Xenopus* hearts, human induced pluripotent stem cells (hiPSCs)-derived cardiac spheroids, and *Caenorhabditis* elegans. Together, these results extend the practical utility of FLFM by enabling robust and efficient volumetric reconstruction across heterogeneous biological specimens and imperfect optical conditions.

## RESULTS

### The principle and framework of HYPER-Net

HYPER-Net establishes a new reconstruction framework for FLFM by combining self-supervised learning with explicit physics conditioning via experimental PSFs. This design enables rapid volumetric inference while improving robustness to optical-system variation and reducing dependence on paired training data. For data acquisition, FLFM employs an epifluorescence platform implemented with a microlens array (MLA) and records multiple parallax views of the specimen in a single camera exposure, thereby encoding volumetric information into a two-dimensional measurement (**Fig. 1a**). Notably, a dual-channel configuration was implemented by integrating spectrally distinct emission filters corresponding to each microlens, enabling simultaneous multicolor fluorescence acquisition^39^.

**Figure 1.**
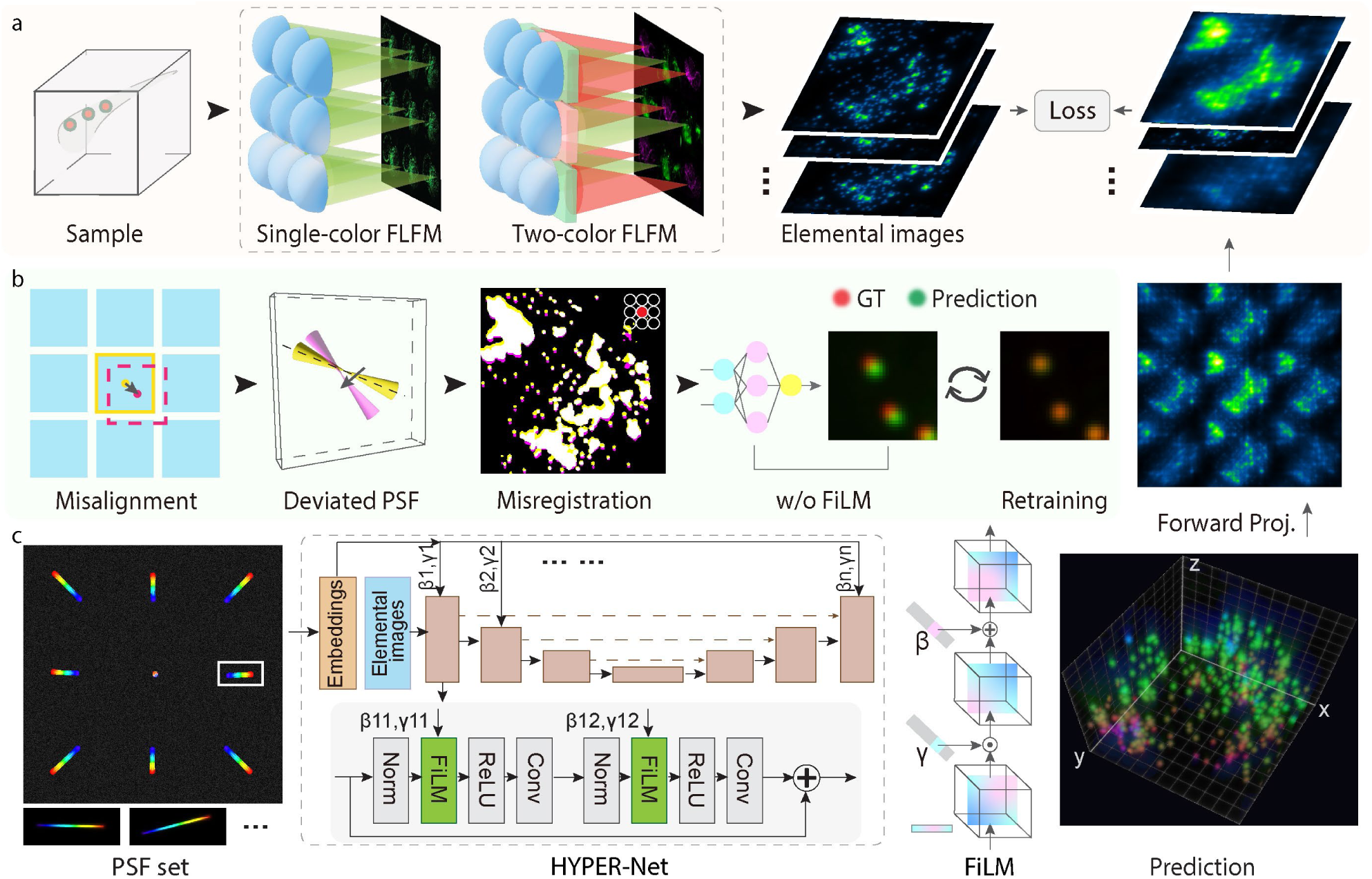
The HYPER-Net principle and architecture. (a) Schematic of Fourier light-field image acquisition. A microlens array (MLA) positioned at the pupil plane encodes the spatial and angular object information into a 3 × 3 array of elemental images on the camera, which serve as the input to the reconstruction network. (b) Variations in alignment and system geometry introduce depth-dependent changes in the elemental point spread functions (PSFs), leading to spatial misregistration and degraded reconstruction fidelity. Such optical deviations alter the forward model relating the reconstructed volume to the measured light-field data and therefore often require retraining in conventional learning-based reconstruction frameworks. (c) Overview of the HYPER-Net workflow. Hybrid PSFs that capture geometry-dependent optical deviations are processed into compact embeddings to generate FiLM parameters (*γ*, *β*), which modulate intermediate feature maps throughout a U-Net backbone. The reconstructed volume is forward-projected through the corresponding elemental PSFs, and the resulting projections are compared with the input elemental images to compute the self-supervised loss. This error is then backpropagated to iteratively update the network parameters during training.

Conventional supervised networks are trained to learn a direct mapping from two-dimensional angular measurements to three-dimensional volumes^32–34^. While effective under matched conditions, these approaches generalize poorly to previously unseen specimen structures or altered optical configurations (**Supplementary Fig. 1**). In addition, variations in the PSFs across experiments often require repeated model retraining (**Fig. 1b and Supplementary Note 1**), thereby limiting robustness, scalability, and practical deployment. In contrast, HYPER-Net employs a physics-guided, self-supervised reconstruction framework (**Fig. 1c and Supplementary Fig. 2**), which extends hybrid PSFs used in conventional Richardson–Lucy (RL) deconvolution^40^ into a learnable framework that adapts the reconstruction to experiment-specific optical conditions. Specifically, the measured high-dimensional PSF stack is processed into a compact embedding vector, which was used to condition feature-wise linear modulation (FiLM^41^) layers distributed throughout a U-Net backbone. Across these layers, intermediate feature maps were adaptively scaled and shifted based on the optical response, thereby maintaining reconstruction conditioning on the system PSF (**Methods and Supplementary Note 2**).

Within the self-supervised framework, the estimated volume is forward-projected through multiple angular PSFs, and the resulting projections are compared with the corresponding measured angular views to compute the loss during training (**Supplementary Fig. 3**). This strategy enforces consistency with the recorded FLFM measurements while allowing PSF-aware modulation to guide feature extraction and volumetric inference. The FiLM parameters adapt intermediate representations to distinct PSF conditions and influence how the decoder maps lateral image cues into axial information. Consistent with this mechanism, the learned FiLM parameters exhibited a strong correlation with the lateral shift slopes of the PSFs at the decoder stages (**Supplementary Fig. 4**), supporting the view that the modulation captures physically meaningful directional characteristics of the image formation process.

This PSF-conditioned strategy substantially improves reconstruction robustness under optical variation. Without such conditioning, PSF-induced geometric mismatch and alignment errors degrade reconstruction quality, resulting in markedly reduced image quality relative to both the PSF-unaware supervised baseline and self-supervised baseline without PSF embedding (**Supplementary Fig. 5**). By contrast, HYPER-Net generates distinct reconstructions under different PSF conditions while maintaining stable outputs for similar or identical PSFs, highlighting both adaptability and consistency across imaging settings. For a reconstructed volume of 460 × 460 × 400 μm^3^, HYPER-Net achieved a 100-fold speed improvement over RL deconvolution. Incorporating the additional PSF-conditioning module, the network still maintained an average inference throughput of 14 volumes per second across 500 trials (**Supplementary Fig. 5**), comparable to the state-of-the-art methods^37^. These results indicate that HYPER-Net offers an efficient and robust solution for volumetric reconstruction under diverse optical conditions.

### Characterization of PSF-conditioned HYPER-Net

We next evaluated HYPER-Net in biological settings to determine whether its PSF-conditioned design confers practical advantages in both reconstruction fidelity and robustness to optical variation. Specifically, we used human colon organoids (HCOs) and beating *Xenopus* embryonic hearts as representative *in vitro* and *in vivo* models to confirm the quality of volumetric reconstruction while preserving sensitivity to experiment-specific PSF conditions.

We first performed Fourier light-field imaging of 5–7-day-old hCOs, which are important models for studying gastrointestinal development, disease, drug absorption, toxicity, and host–microbiome interaction^42,43^. At this developmental stage, the organoids exhibited substantial morphological heterogeneity, providing a stringent test for volumetric reconstruction (**Fig. 2a**). HYPER-Net yielded higher spatial resolution and improved signal-to-background ratio than conventional RL deconvolution (**Figs. 2b, c, and Supplementary Fig. 6**), while reducing reconstruction time by approximately 100-fold relative to RL deconvolution with 10 iterations. Notably, HYPER-Net also outperformed axial wide-field scanning stacks, benefiting from the substantially extended depth of focus (approximately 375 times) using FLFM, which alleviates the severe out-of-focus blur that commonly affects thick biological specimens.

**Figure 2.**
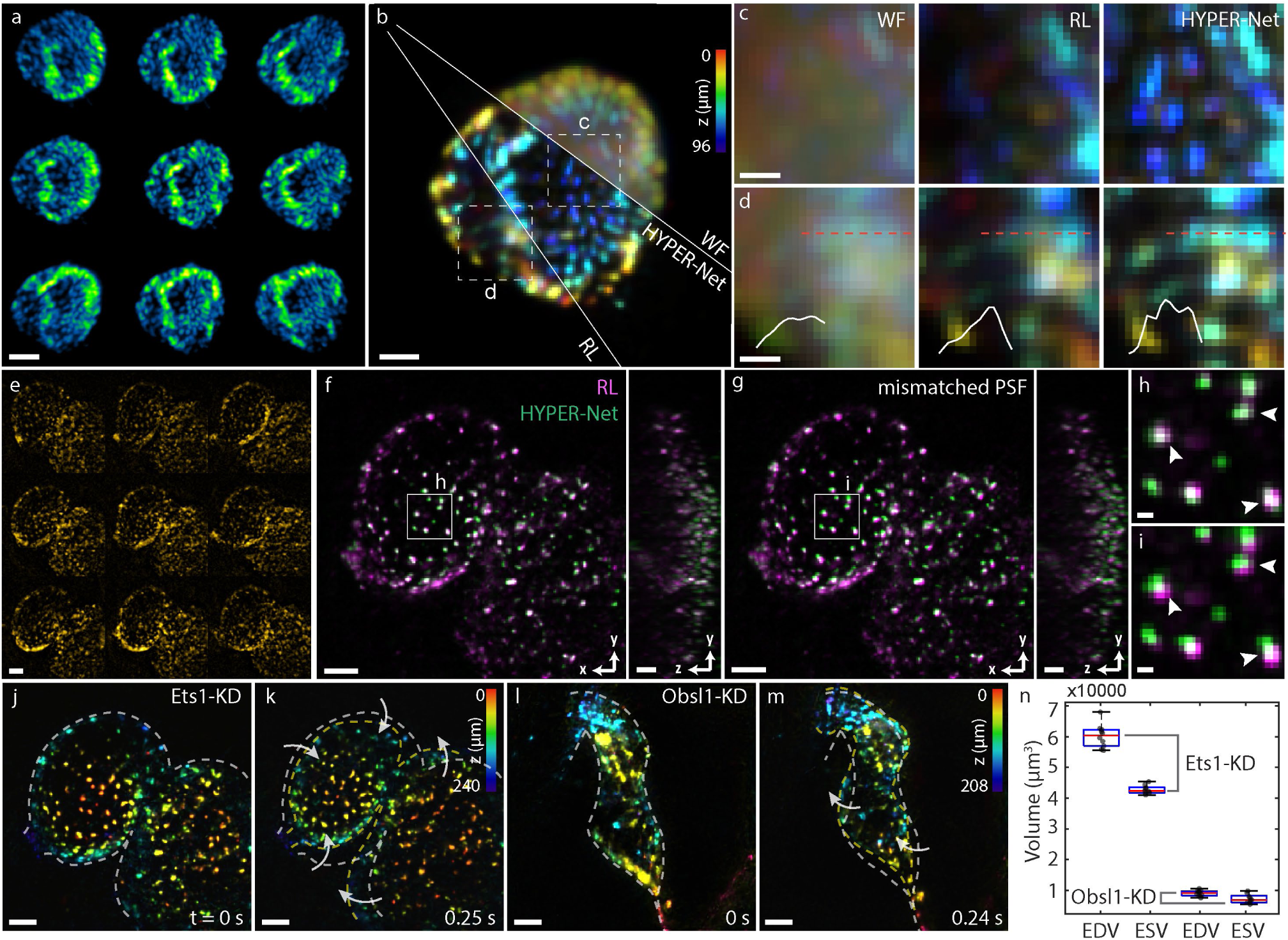
Characterization of PSF-conditioned HYPER-Net. (a) Raw elemental images of human colon organoids (hCOs) stained for nuclei with SYTO16. (b) Corresponding depth-color-coded volumetric reconstructions acquired using scanning wide-field (WF) images, conventional Richardson-Lucy (RL) deconvolution, and HYPER-Net. (c, d) Zoomed-in images of the boxed regions as marked in (b), showing the improved contrast and spatial resolution achieved by HYPER-Net. (e) Raw elemental images of H2B: GFP-labeled cardiomyocyte nuclei in an embryonic *Xenopus* heart. (f) Corresponding three-dimensional maximum-intensity projection (3DMIP) of the volume reconstructed using RL deconvolution (magenta) and HYPER-Net (green). (g) Comparative HYPER-Net reconstruction with an intentionally mismatched PSF condition, implying the role of FiLM modulation. (h, i) Zoomed-in images of the boxed regions in (f, g), showing that the mismatched condition leads to clear spatial discrepancies in the reconstructed nuclei. (j-m) Depth-color-coded volumetric reconstructions of embryonic *Xenopus* hearts using HYPER-Net at systole and diastole for *Ets1*-KD (j, k) and *Obsl1*-KD (l, m) tadpoles, revealing impaired cardiac pumping in the knockdown hearts. (n) Box plots of end-diastolic ventricular volume (EDV) and end-systolic ventricular volume (ESV) measured from the *Ets1*-KD heart (j, k) and the *Obsl1*-KD heart (l, m) over ten cardiac cycles. Scale bars: 50 μm (a, b, f, g, j-m), 10 μm (c, d, h, i), 20 μm (e).

We next applied HYPER-Net to the volumetric analysis of cardiac development in gene-edited *Xenopus* laevis embryos. *Xenopu*s provides an enabling vertebrate model for studying cardiac morphogenesis because its early embryonic heart is readily accessible, developmentally well-defined, and highly amenable to genetic perturbation^44,45^. The scan-free volumetric acquisition of FLFM enabled efficient imaging of beating hearts over extended cardiac cycles, covering a field of view of approximately 500 µm laterally and 200–300 µm axially at the targeted developmental stage (**Fig. 2d**). However, practical use of FLFM in this context remains limited by reconstruction throughput and by the requirement for accurate PSF modeling, as PSF mismatch can introduce spatial misregistration and attenuation of true biological signals (**Supplementary Fig. 7**).

Because HYPER-Net explicitly conditions reconstruction on the input PSF, we tested whether this conditioning directly affects the reconstruction output. Indeed, different PSF conditions produced distinct reconstructed volumes. With the PSF matched to the experimental configuration, HYPER-Net generated a volume consistent with the reference RL reconstruction. By contrast, conditioning on a mismatched PSF resulted in clear three-dimensional rotation and spatial misalignment relative to the reference volume (**Figs. 2e-h**). We then used this physics-informed framework to evaluate phenotypic consequences of genetic perturbation during cardiac development. The *Ets1*-mutant heart exhibited largely preserved gross morphology but reduced cardiac output (**Fig. 2i, j, Supplementary Movie 1**), consistent with impaired functional development and extending previous reports^46^. By contrast, the *Obsl1*-mutant heart displayed a narrowed and shortened ventricular chamber with minimal volumetric change during systole (**Fig. 2k, l, Supplementary Movie 2**), indicative of severely compromised contractility and reduced pumping capacity (**Fig. 2m**)^47^. These findings are consistent with an essential role of *Obsl1* in maintaining sarcomeric organization and mechanical stability in muscle cells^48^. Overall, these results validate HYPER-Net as a robust platform for high-throughput volumetric phenotyping and support its utility for probing genetic mechanisms underlying congenital heart disease.

### Imaging hiPSC-derived cardiomyocytes with HYPER-Net

Three-dimensional microtissues formed from hiPSC-derived cardiomyocytes (hiPSC-CMs) recapitulate key structural and functional features of the human heart and therefore serve as valuable *in vitro* models for studies of cardiac physiology, disease, regeneration, and pharmacological toxicology^49–51^. Light-field imaging enables ultrafast volumetric visualization of both subcellular motion and calcium signaling throughout intact cardiac spheroids^46,52^. We therefore used this system to evaluate HYPER-Net in a particularly demanding regime, where structural motion and functional calcium activity are tightly coupled in space and time. Specifically, we imaged 3D hiPSC-CM spheroids labeled with a ratiometric calcium indicator (**Fig. 3a**). Ratiometric calcium imaging is especially advantageous for cardiac microscopy because the signal is less sensitive to variations in dye loading, photobleaching, and changes in cell position or volume^53^. Using the dual-channel configuration, rhythmic fluorescence fluctuations from the two spectra were recorded simultaneously (**Fig. 3b**). The time-lapse observation revealed calcium activity, with red emission predominating at low calcium levels and green emission increasing as calcium concentration rose (**Fig. 3c-f**).

**Figure 3.**
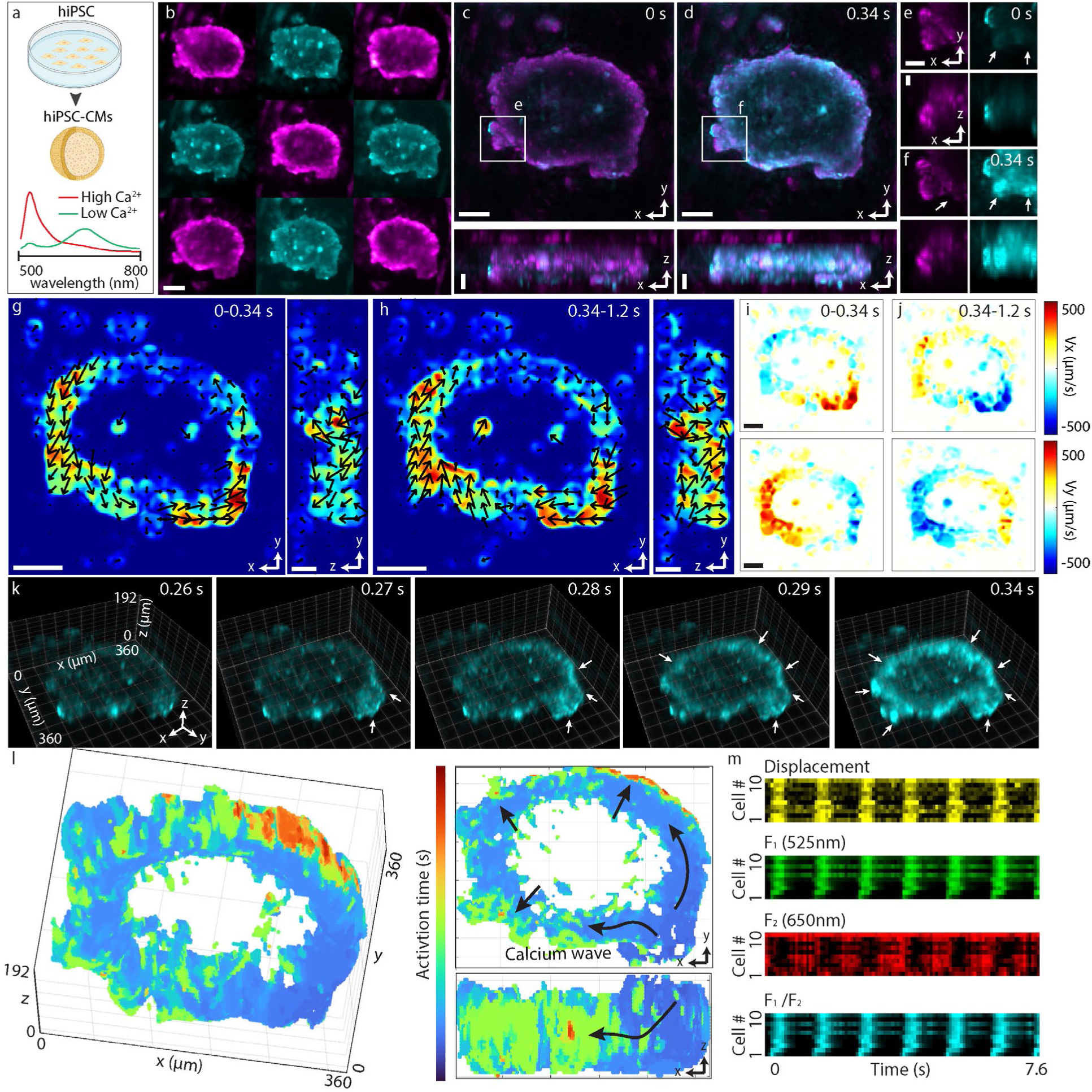
Imaging hiPSC-derived cardiomyocytes with HYPER-Net. (a) Schematic of hiPSC-CM spheroids and the fluorescence response of the ratiometric calcium indicator, showing opposite signal changes in the high- and low-Ca^2+^ channels with changing intracellular calcium. (b) Raw two-color elemental images from 3D hiPSC-CM spheroid labeled with Calbryte Red 525/650 AM. (c, d) Two-color 3DMIPs of the HYPER-Net-reconstructed spheroid during relaxation (c, *t* = 0 s) and contraction (d, *t* = 0.34 s). Elevated intracellular calcium is associated with increased 525 nm emission (cyan) and decreased 650 nm emission (magenta). (e, f) Zoomed-in images of the boxed regions in (c) and (d), respectively. Arrows indicate regions showing opposite channel responses during contraction, with increased intensity in the 525-nm channel and decreased intensity in the 650-nm channel at high calcium levels. (g, h) Displacement magnitude maps during contraction (g) and relaxation (h) over one beating cycle. Arrows indicate the local direction of spheroid motion, which reverses between contraction and relaxation. (i, j) Velocity maps along the x- and y-axes during contraction (i) and relaxation (j), showing inward contractile motion followed by outward relaxation. (k) Time-resolved 3D views of the 525-nm channel from 0.26 to 0.34 s, showing propagation of calcium activity at the onset of contraction. Arrows indicate regions of elevated calcium activity. (l) Three-dimensional activation map of calcium propagation across the spheroid shell. Insets show corresponding two-dimensional projections. (m) Heatmaps of displacement, 525-nm fluorescence, 650-nm fluorescence, and ratiometric calcium signal for 10 tracked cells over time. Scale bars: 100 µm (b, g, h), 50 µm (c, d, i, j), 20 µm (e, f).

The high volumetric acquisition rate of 100 Hz, combined with low epifluorescence excitation (average excitation power <0.4 mW over a 1 mm × 1 mm field at 100 Hz), enabled recording for thousands of frames without detectable photodamage (**Supplementary Fig. 8, Supplementary Movies 3, 4**). These datasets captured both the morphological deformation of the spheroids during the contraction cycle (**Fig. 3g-j**) and the associated calcium flux occurring on a timescale of tens of milliseconds (**Fig. 3k**). From the reconstructed volumes, we derived three-dimensional activation maps that revealed a clear spatiotemporal propagation pattern of calcium signaling, initiated from the one region of the spheroids and extending from the inner core toward the outer shell (**Fig. 3l**). Notably, such three-dimensional maps provide direct insight into conduction pathways within the tissue and may enable localization of arrhythmogenic initiation sites^54^.

Furthermore, the reconstructed volumes permitted tracking of individual cells throughout repeated beating cycles (**Supplementary Note 3**), allowing simultaneous quantification of cell motion trajectories and calcium transients across the entire spheroids (**Fig. 3m**). The resulting cyclic displacement and fluorescence dynamics define phase-space trajectories for individual cardiomyocytes, revealing a positive association between structural motion and calcium-driven functional activity (**Supplementary Fig. 8**). Since HYPER-Net is self-supervised and explicitly conditioned on the optical forward model, the framework reduces dependence on paired experimental ground-truth volumes and generalizes across specimens with diverse biological architectures. The results exhibited comparable quality to conventional RL deconvolution while offering substantially enhanced computational efficiency, with more than 200-fold faster processing for complex samples and a reduction in runtime from 6 hours to 1 minute for a 15 s sequence. We expect the improvement to substantially decrease dependence on experimentally curated reconstruction pipelines and enable high-throughput, time-resolved volumetric interrogation of cardiac excitation-contraction coupling across diverse physiological and pathological conditions.

### Study of locomotion-associated neural activity in freely moving *C*. *elegans*

Finally, we evaluated HYPER-Net for volumetric calcium imaging in freely moving *Caenorhabditis elegans* by recording activity in body-wall muscles and motor neurons during unconstrained locomotion. *C. elegans* is a well-established model for studying neural circuit function because of its optical transparency, compact nervous system, fully mapped cell lineage, and complete neuronal connectome^55,56^. However, conventional two-dimensional imaging often fails to maintain focus during its three-dimensional motions, and therefore typically requires physical confinement under coverslips or in microfluidic devices, potentially perturbing natural physiological states. In contrast, the FLFM platform enabled imaging of freely swimming worms at both larval (**Fig. 4a-c**) and adult stages (**Fig. 4d-m**) across a near-millimeter lateral field of view and more than 100 µm axial depth at 100 volumes per sec. To monitor body-wall muscle activity, we used GCaMP3 as a calcium-sensitive reporter together with calcium-insensitive RFP as a reference^57^ (**Fig. 4a**). Muscles on the actively bending side exhibited elevated calcium activity during contraction (**Fig. 4b**). The progressive shift in calcium activity along the body over time further revealed propagation of the bending wave, for example, during tail-bend initiation (**Fig. 4c**). These results demonstrate that HYPER-Net can resolve fast, distributed muscle activation patterns during natural three-dimensional locomotion.

**Figure 4.**
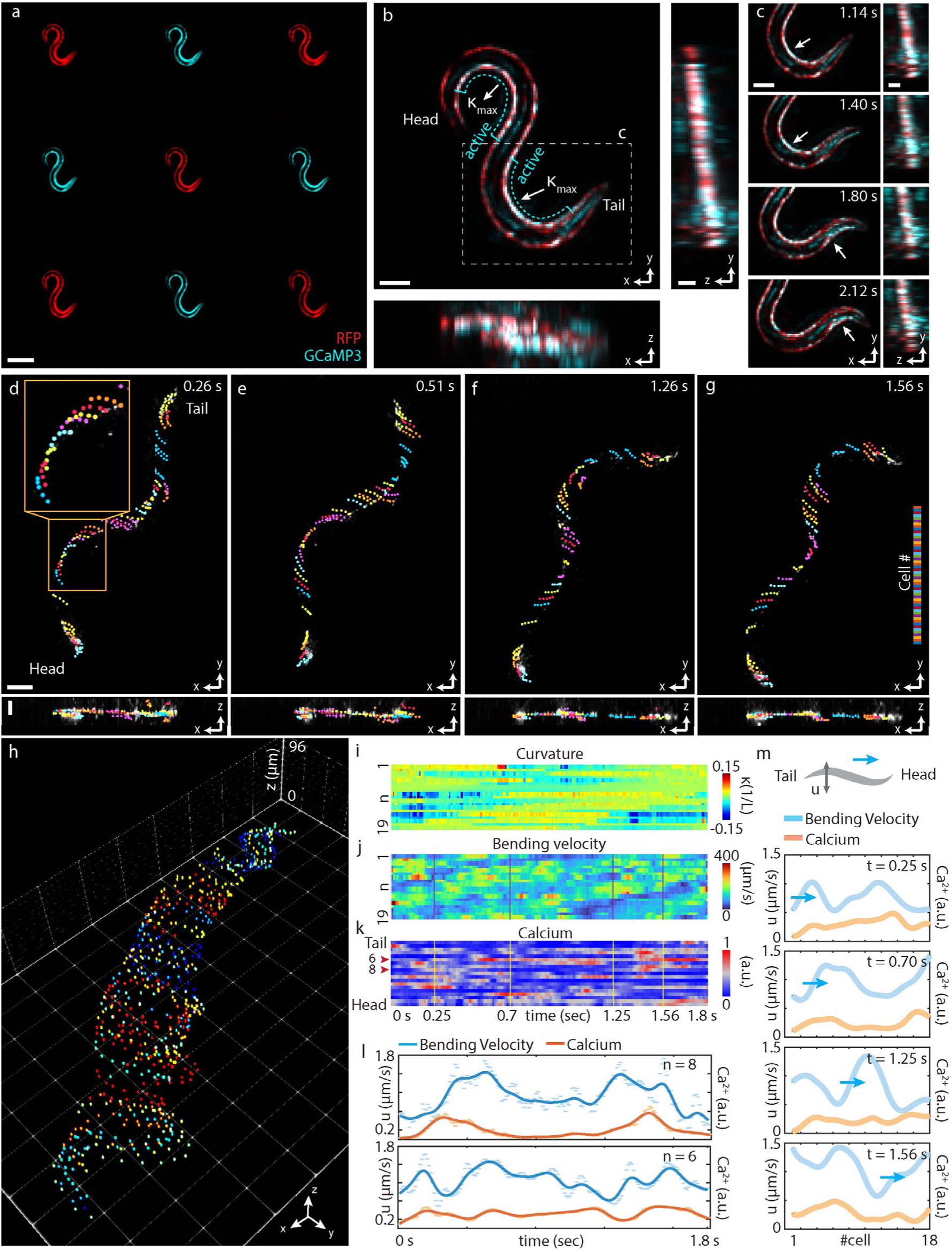
Imaging locomotion-associated neural activity in freely moving *C. elegans* with HYPER-Net. (a) Raw two-color elemental images of a *C. elegans* larva expressing GCaMP6s (cyan) and a calcium-insensitive RFP reference (red) in body-wall muscles. (b) Two-color 3DMIP of the reconstructed larval volume. (c) Time-lapse images from the boxed region in (b) at 1.14, 1.40, 1.80, and 2.12 s, showing increasing calcium activity in active body-wall muscles (white arrows) during tail-bend initiation. (d-g) Two-color 3DMIPs of reconstructed volumes from an adult *C. elegans* expressing GCaMP6s and mNeptune in motor neurons, shown at 0.26, 0.51, 1.26, and 1.56 s. Colored dots indicate tracked single motor-neuron positions, and traces show trajectories across five successive frames separated by 0.05 s. (h) Overlay of motor-neuron trajectories over 1.81 s at 0.1 s intervals, revealing coordinated whole-body undulatory motion. (i-k) Heatmaps of 3D body curvature, lateral bending velocity, and motor-neuron calcium activity over time, showing periodicity in curvature and bending velocity and a correspondence between calcium activity and bending velocity. (l) Representative calcium signals and corresponding bending velocities for two tracked motor neurons, showing temporally correlated peaks. (m) Calcium activity and bending velocity for all tracked motor neurons, ordered from tail to head, at the time points marked in (j) and (k), illustrating propagation of bending waves during locomotion. Scale bars: 100 µm (a, d top), 25 µm (b, c, d bottom).

We next examined motor-neuron-associated activity in freely moving adult worms labeled with calcium indicators, which captured the rhythmic neural dynamics underlying coherent locomotor behaviors such as crawling and swimming^58^. High-fidelity sequential volumetric reconstructions of the mNeptune channel enabled accurate tracking of neuronal cell bodies at single-cell resolution despite rapid three-dimensional body motion (**Fig. 4d-h, Supplementary Fig. 9, Supplementary Note 4**). The time-lapse results also revealed rhythmic bending across successive body segments, generating a head-to-tail curvature wave during forward locomotion (**Fig. 4i, Supplementary Movies 5, 6**). Notably, bending velocity reached its maximum as the body passed through a straightened configuration while reversing bending direction from one side to the other (**Fig. 4j**). Correspondingly, calcium activity in motor neurons derived from the GCaMP6s channel (**Fig. 4k**) increased with bending speed **(Fig. 4l**) and formed a spatiotemporal wave propagating along the elongated body during locomotion (**Fig. 4m**). These results verify HYPER-Net as a practical framework for high-speed volumetric interrogation of locomotion-linked neural and muscular dynamics in freely behaving animals.

## DISCUSSION

Advances in biological volumetric imaging are increasingly driven by the joint design of optical encoding and computational reconstruction, enabling image quality and imaging speed beyond what either strategy can achieve alone^59^. Light-field imaging offers an efficient route to single-shot three-dimensional acquisition, but its broader applicability has been limited by the computational burden and reconstruction artifacts associated with iterative model-based approaches, as well as by the restricted generalizability of supervised learning methods to specimen variation and optical mismatch. Here, we address these limitations by introducing HYPER-Net, a physics-conditioned, self-supervised reconstruction framework that combines rapid volumetric inference with improved robustness to experiment-specific PSF variation, thereby enabling practical volumetric reconstruction across diverse biological settings. HYPER-Net advances light-field reconstruction by dual use of the PSF prior: as the forward operator that enforces measurement consistency and as a conditioning signal that modulates intermediate feature representations via FiLM (**Methods**). This integrated design reduces sensitivity to optical deviations and improves reconstruction stability under changing imaging conditions. In the present implementation, FiLM parameters are generated from PSF lateral-shift slopes using multilayer perceptrons, yielding an efficient, computationally tractable strategy for PSF-aware conditioning. The framework is readily extensible to more expressive architectures, including residual, attention-based, and CNN–transformer backbones^60,61^ (**Supplementary Note 2**), and is compatible with emerging hardware configurations such as adaptive optics and metasurface-enabled imaging platforms^62,63^. The capabilities of HYPER-Net were demonstrated across multiple biological models spanning distinct morphologies, motion regimes, and fluorescence labeling strategies, including organoids, embryonic hearts, cardiac spheroids, and freely moving *C. elegans*. By enabling fast, high-fidelity, and robust volumetric reconstruction, HYPER-Net supports downstream interrogation of structural and functional dynamics in complex living systems. We anticipate that this framework will facilitate high-throughput and long-term volumetric imaging across applications such as cardiac functional assessment^64^, pharmacological evaluation^65^, developmental phenotyping^66^, and systems neuroscience^67^, and will serve as a broadly useful paradigm for integrating optical physics with machine intelligence in computational microscopy.

## METHODS

### Experimental setup

The study was conducted on the home-built Fourier light-field microscopy system, capable of single-volume and simultaneous dual-color volumetric acquisition^39,68^. The system was equipped with a water-dipping physiology objective lens (CFI75 LWD 16×/0.8NA W, Nikon) for a long working distance and large imaging field of view. For single-color imaging, samples were epi-illuminated by a single-color cold visible mounted light-emitting diode (LED) peaked at 470 nm (M470L5, Thorlabs), which was filtered by an excitation filter (MF469-35, Chroma) and was reflected by a dichroic mirror (89402bs, 50.8 × 72 × 2mm, Chroma). Two achromatic doublets (AC254–200-A-ML and AC254–400-A-ML, Thorlabs) were inserted between the objective lens and the LED to form Kohler illumination. The fluorescence emitted from the sample was collected by the objective lens and was filtered by the dichroic mirror and an emission filter (89402 m, 50 mm diam, Chroma) and was focused as a wide-field image by a 300-mm tube lens (AC508–300-A-ML, Thorlabs) on the native image plane (NIP). The Fourier light-field modality consisted of a 200-mm Fourier lens (AC508–200-A-ML, Thorlabs) placed with its front focal plane on the NIP, and a customized microlens array (MLA) assembled by inserting 3-by-3 planoconvex lenses into a lens mount. Each 3.3-mm square lens was diced from a commercially available planoconvex lens with a 20-mm diameter and a 30-mm focal length (#45-240, Edmund Optics). Videos were recorded with an sCMOS camera (ORCA-Flash-4.0 v3, Hamamatsu) using the central 1536 × 1536 pixels to capture all elemental images at 100 Hz. For the two-color imaging scheme, samples were illuminated simultaneously using light sources peaked at 470 nm (M470L5, Thorlabs) or 590 nm (M590L4, Thorlabs), both filtered with ZET594/10× (Chroma). On the back focal plane of the Fourier lens, the microlens array was assembled with 9 customized microfilters (Chroma, 3.3 mm × 3.3 mm × 1 mm, ET520/40m for the GFP channel, ET632/60m for the mCherry channel) to enable simultaneous multi-color acquisition.

### Pipeline of HYPER-Net

To enable physics-aware volumetric reconstruction from FLFM measurements, we designed a point-spread-function (PSF)-conditioned deep neural network that integrates optical system parameters via feature-wise linear modulation. The elemental images from different angular views were concatenated along the channel dimension to form the network input. Optical system characteristics were represented by an 18-dimensional vector describing depth-dependent PSF parameters (lateral shift coefficients in *x* and *y*). These parameters were encoded into a compact embedding vector using a multilayer perceptron with four fully connected layers and LeakyReLU activations, providing global conditioning for reconstruction. The reconstruction backbone followed a 2D U-Net architecture, an effective approach for capturing multi-scale spatial context while preserving fine structural details. The network employed a four-level encoder-decoder structure with residual blocks, each comprising GroupNorm, FiLM modulation, LeakyReLU activation, and 3 × 3 convolutional layers. PSF information was incorporated at every block via FiLM conditioning, where channel-wise scaling and shifting parameters were predicted from the PSF embedding:

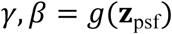

where *g*(⋅) denotes a linear projection generating 2*C* parameters for a feature map with *C* channels. Given normalized feature maps *h*, FiLM modulation was applied as:

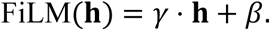

The network predicted a volumetric output, represented as channel-wise slices, after a prediction head consisting of sequential 3 × 3 convolutional layers with LeakyReLU activations. Finally, the self-supervised module performs forward projections of the estimated volume through multiple angular PSFs, minimizing the loss between the projections and the corresponding angular measurements in each iteration. To simulate depth-dependent variations in the PSF lateral shift due to optical aberrations and mechanical misalignment, we generated 100 hybrid point spread functions (hPSFs) using randomly perturbed microlens array configurations (**Supplementary Fig. 10**). Individual microlenses were displaced by up to 100 µm, and the entire microlens array holder was shifted by up to 1 mm. These perturbation magnitudes were selected to reflect the fabrication tolerances of the 3D-printed mounting structure and the alignment deviations observed under practical experimental conditions. For each perturbed configuration, rayoptics simulations were performed to compute the corresponding depth-dependent lateral shift slopes in the *x–z* and *y–z* planes. These slopes defined continuous lateral-shift trajectories of PSF locations across axial depth in different elemental images. A wave-optics–simulated PSF intensity profile was subsequently translated along the fitted trajectories to construct the final hybrid PSF (**Supplementary Notes 2, 5**).

### Preparation and staining of hiPSC-derived colon organoids

The frozen vial of human iPSC-derived colon organoids (hCOs) was purchased from Millipore Sigma (SCC 300). Organoid fragments were embedded in 25-μL growth factor-reduced Matrigel (GFR MG, Corning 356231) domes (protein concentration 8 mg/mL). After the gels solidified in the incubator, 500-750 μL of media was added. For the first 3 days after thawing or passaging, ROCK inhibitor Y-27632 (R&D Systems 1254) was added to the media to prevent cell death. Then the media were exchanged every 2 days until the following passage. Organoids used in this work are generally at day 5–7 after passaging^42^.

### Preparation, staining, and imaging of 3D hiPSC-CMs

MR90-IV (WiCell Research Institute) was maintained in mTeSR1 medium (Stem Cell Technologies) and induced for cardiac differentiation as previously described^69^. Briefly, hiPSCs were treated with 100 ng/ml recombinant human activin A (R&D Systems) in RPMI medium with 2% B27 insulin-free (RPMI/B27 insulin-free medium) on day 0. After 24 h, the medium was replaced with 10 ng/ml recombinant human bone morphogenetic protein-4 (BMP4, R&D Systems) in RPMI/B27 insulin-free medium, as used from days 1 to 4. The medium was changed to RPMI medium with 2% B27 containing insulin (RPMI/B27 medium) on day 4.

Cardiac spheroids were generated on day 10 post-differentiation. Differentiated cells were dissociated with 0.25% trypsin/EDTA and seeded into AggreWell 400 plates (#34415, Stem Cell Technologies) at 1,500 cells/microwell to allow cells to form cardiac spheroids. Before cell seeding, plates with 1 ml/well of RPMI/B27 medium were centrifuged at 1000 g to release trapped bubbles in microwells. To prevent cell death, the medium was supplemented with 10 µM of Rock inhibitor Y-27632 (Selleck Chemicals). Plates were centrifuged at 100 g to distribute the cells and then placed in an incubator. After 24 h, spheroids were harvested to remove the Rock inhibitor, and the RPMI/B27 medium was replaced in the suspension culture. 3D spheroids were maintained until the day of testing (< 30 days post-differentiation). The medium was refreshed every 4-5 days.

On the day of imaging, 9.2 μL anhydrous dimethyl sulfoxide (DMSO, ThermoFisher Scientific, D12345) was added into one vial of Cal RedTM R525/650 potassium salt (AAT Bioquest, 20588) to make a vial stock solution and 2.3 μL stock solution was diluted into 2.3 ml of prewarmed complete OptiMEM (OptiMEM, ThermoFisher Scientific, 51985-034, with 1% GlutaMAX, ThermoFisher Scientific, 35050-061; 1% NEAA, ThermoFisher Scientific, 11140-050; and AlbuMAX 2 g/L, ThermoFisher Scientific, 11020-021) to make a loading solution at a final concentration of 5 μM. Then, the original maturation medium was replaced with fresh culture medium containing Cal RedTM R525/650, and the cells were incubated in the incubator (37 ℃, 5% CO_2_) for 85 min. After that, the staining medium was aspirated. Cells were recovered in a 3 ml prewarmed maturation medium for 20 min. In the end, the maturation medium was replaced with complete OptiMEM, and the dish containing cells was placed on a coffee warmer under the microscope to maintain the medium temperature during imaging. A full-frame video was recorded at 100 Hz, lasting over 15s for each 3D hiPSC-CM, covering more than 7 beating cycles.

### Preparation and imaging of *Xenopus laevis*

*Xenopus laevis* embryos were obtained by fertilizing wild-type oocytes with wild-type testis (ordered from Xenopus 1 Corp). The fertilized embryos were micro-injected at the 4-cell stage bilaterally with 0.5 ng H2b-GFP RNA (1 ng RNA per embryo) in their dorsal marginal zone (prospective cardiac region). 7.5 ng translation-blocking MO of Ets1 (ordered from Gene Tools) was also injected bilaterally to mute the ETS1 gene. The fertilized embryos were screened for fluorescent signals in the heart and raised to the mature tadpole stage (stage 46+) as described^70^.

On the day of the experiment, embryos were placed ventral side up on a 3D printed bed in 0.1× MMR plus 0.1% MS222, and a video of 10 s (∼14 cardiac cycles) was taken at 100 Hz for each tadpole. All experimental procedures were performed according to the 429 USDA Animal Welfare Act Regulations and were approved by the Institutional Animal Care and Use Committee in compliance with Public Health Service Policy.

### Preparation and imaging of *C. elegans*

Strains were maintained at 20℃ and were fed E. coli (OP-50) bacteria on agar plates using standard protocols. The transgenic line AQ2953 ljIs131[Pmyo-3::GCaMP3-SL2-tagRFP-T] was used to perform calcium imaging on muscle cells^71^. The strain ZM9624 (lin-15(n765) X; hpIs675) hpIs675 [rgef-1p::GCaMP6s::3xNLS::mNeptune + lin-15(+)] expresses GCaMP6s and mNeptune panneuronally in motor neurons. All worms were cultured on standard nematode growth medium plates seeded with OP50 and maintained in incubators at 20°C for about 72 hours after hatching. On the day of imaging, 3% (w/w) Methylcellulose was prepared by dissolving Methylcellulose in S. Basal. Worms were picked into a droplet of solution, sandwiched between two glass slides separated by No. 1 glass coverslips (0.15 mm).

## DATA AVAILABILITY

All data needed to evaluate the conclusions in the paper are present in the paper and/or available upon request from the corresponding author.

## CODE AVAILABILITY

The code for HYPER-Net is available as Supplementary Software. The code has been written in MATLAB (MathWorks) and tested with version 2022b. To install the package, unzip the compressed folder and follow the instructions in the file readme.txt. The latest version of the software will be available at https://github.com/ShuJiaLab/HYPER upon publication.

## ACKNOWLEDGEMENTS

We acknowledge the support of the National Institutes of Health grants R35GM124846 (to S.J.), R21HD110918 (to S.N. and S.J.), and R01AA028527 (to C.X.), and the National Science Foundation grants BIO2145235 (to S.J.) and CBET2225990 (to S.J. and S.N.).

## AUTHOR CONTRIBUTIONS STATEMENT

Z.L. and S.J. conceived and designed the project. Z.L. and X.H. developed the algorithm. Z.L. and P.C. helped with the coding environment. Z.L. and W.L. performed imaging experiments. G.K prepared the colon organoid sample. P.F. prepared the cardiac spheroid sample. J.H. prepared the frog embryo sample. L.P. prepared the *C. elegans* sample. P.C. helped interpret the *C. elegans* results. H.W. and Z.L. conducted image analysis and generated results. S.T., H.L., S.N., and C.X. inspected the research results. S.J. supervised the overall project. Z.L. and S.J. wrote the manuscript with input from all authors.

## COMPETING INTERESTS STATEMENT

The authors declare no competing interests.

## Notes

### Competing Interest Statement

The authors have declared no competing interest.

